# Persistent neural activity encoding real-time presence of visual stimuli decays along the ventral stream

**DOI:** 10.1101/088021

**Authors:** Edden M. Gerber, Tal Golan, Robert T. Knight, Leon Y. Deouell

## Abstract

Neural populations that encode sensory experience should be persistently active for as long as the experience persists. However, research into visual neural activity has focused almost exclusively on onset-driven responses that cannot account for sustained perception. We used intracranial recordings in humans to determine the degree to which the presence of a visual stimulus is persistently encoded by neural activity. The correspondence between stimulus duration and response duration was strongest in early visual cortex and gradually diminished along the visual hierarchy, such that is was weakest in inferior-temporal category-selective regions. A similar posterior-anterior gradient was found within inferior temporal face-selective regions, with posterior but not anterior sites showing persistent face-selective activity. The results suggest that regions that appear uniform in terms of their category selectivity are dissociated by how they temporally represent a stimulus in support of real-time experience, and delineate a large-scale organizing principle of the ventral visual stream.

## Introduction

Although visual perception of scenes or objects typically occurs over an extended time window, what we know of human high level visual perception is largely based on onset responses, that is, responses to change (e.g. the EEG/intracranial face-selective N170/N200; Bentin et al. 1996; Allison et al. 1999). While onset responses hold a wealth of information on the functional properties of the respective neural population, they cannot disentangle the transient response to change, from neural activity driven by the ongoing presence of a percept. Thus, how real-time experience persists beyond the onset is unknown, leaving the neural basis of the brunt of perceptual time unaccounted for. Put simply, our question is how do we know, in real-time and beyond the initial detection, whether a stimulus we gaze at is still present 500 or 1500 ms later, and how do we know its content, e.g. whether it is (still) a face or an object? This question is distinct from that of explicit duration estimation tasks (e.g. was a probe stimulus shorter or longer than a reference stimulus; Buhusi & Meck 2005; Lewis & Miall 2003) in which judgments are performed post-hoc, that is, after stimulus offset. Here, we are concerned with how the visual system codes the continued presence of an object in real time. Some evidence of sustained responses to visual stimuli can be gleaned from single unit (Kulikowski et al. 1979; Petersen et al. 1988; Ikeda & Wright 1974) or fMRI studies (Gilaie-Dotan et al. 2008) using long duration visual stimuli, and from variable duration EEG\MEG responses in an explicit duration estimation task (N’Diaye et al. 2004; Pouthas et al. 2000), however these studies did not directly examine the neural basis of sustained, real-time perception.

We recorded from ten human subjects implanted with subdural electrodes over visual cortices using electrocorticography (ECoG) while subjects were engaged in a novel target detection paradigm, with faces and objects presented for variable durations. Since even a brief stimulus can produce a sustained response (Fisch et al. 2009), using variable stimulus durations was crucial for isolating the part of the response driven specifically by the stimulus ongoing presence. However, no duration estimation was required.

We found that early visual cortex (EVC) high frequency broadband response closely tracked the time course of the stimulus, and this precision decreased along the visual hierarchy, i.e. from V1/V2 to V3/V4, from early visual cortex to inferior temporal (IT) cortex, and from posterior to anterior IT. Only the posterior part of IT robustly encoded the presence of the stimulus over time. This posterior-anterior gradient could not be explained by signal-to-noise ratio or by differential influences of attention or saccade-related activity.

## Results

Subjects viewed grayscale images of faces, objects, or other miscellaneous images presented for a duration of 300, 600, 900, 1200 or 1500 milliseconds. To maintain attention subjects responded via button press to a rare target category (clothes, Fig. 1A). All subjects detected the targets successfully (mean±SD hit rate 90.4%±8.6%, false alarm rate 1.9%±3.2%). Responses to the rare targets were not included in the following analyses. Stimulus-induced modulation of the ECoG signal was measured as an increase in the power of high frequency broadband signal (HFB, >30 Hz), which was shown to correlate with local neural firing rate (Manning et al. 2009). Fig. 2 shows characteristic single-trial and average responses. Out of 1067 electrodes, subsequent analysis was performed on 292 electrodes designated as visually-responsive, based on a significant modulation of HFB power to at least one stimulus category.

**Figure 1:**
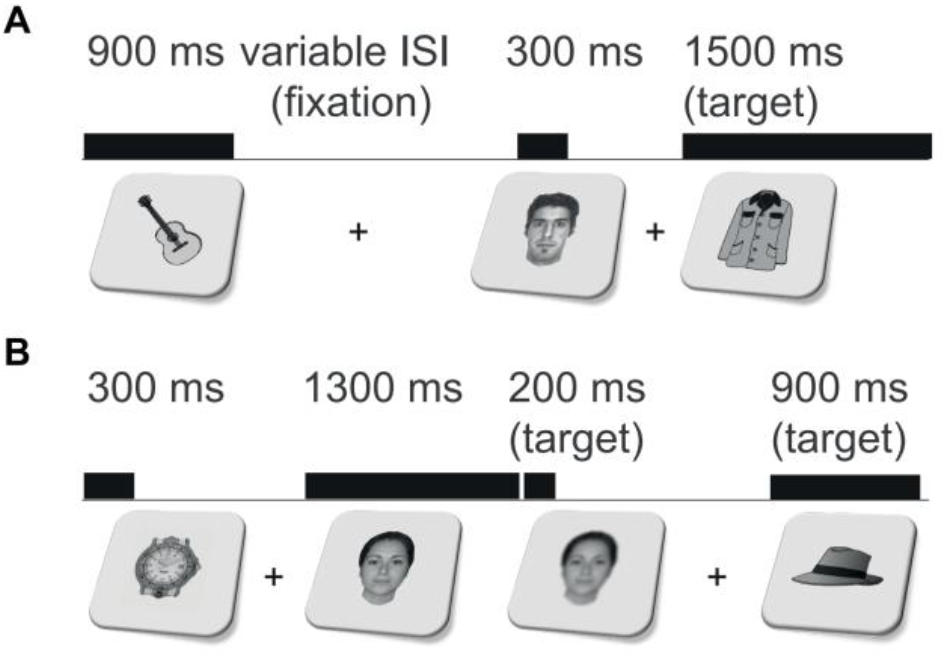
Experimental paradigm. (**A**) Images were presented for 300, 900, or 1500 ms (for 3 subjects also 600 or 1200 ms) with variable inter-stimulus interval (ISI) during which a fixation cross was presented. Subjects responded with a button press to presentation of targets (clothing images; 10% of trials). (**B**) Dual-task control. This task was identical to the first except that subjects also had to respond to rare blurring of the image in the last 200 ms of its presentation.

**Figure 2:**
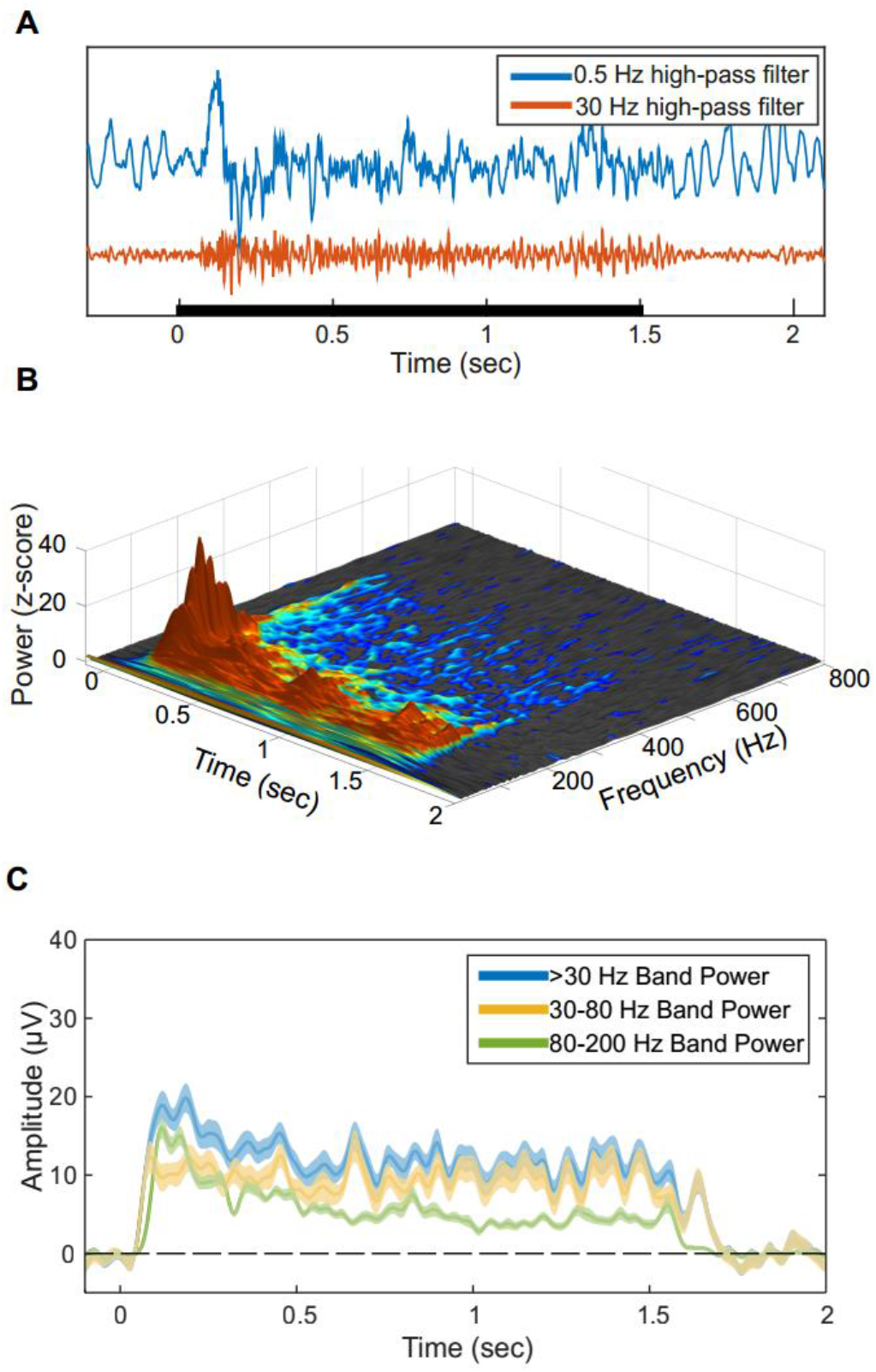
Representative spectral response profile. (**A**) The high-frequency response is evident in raw single trials. Blue and Red traces correspond to 0.5 Hz and 30 Hz high-pass filtered traces respectively of a single trial from an early visual cortex (V1/V2) electrode. Black bar indicates the stimulus duration. (**B**) Time-frequency response to 1500 millisecond visual stimuli averaged across both categories, in the same electrode. Z-axis and color scale represent z-score of the power at each time-frequency data point as normalized by the baseline for the same frequency band. Since both the baseline and response spectra are dominated by a 1/f power law in terms of mean power and variance, this normalization makes it possible to observe relative modulations in high frequency bands which are otherwise too small in comparison to lower frequencies. Color and grayscale data points represent significant and non-significant z-score values respectively (p<0.05, uncorrected). Note that while the early response components contain higher power at high frequencies, the upper bound of significantly above-baseline activity is roughly fixed at 400-500Hz throughout the activity time course, implying a general attenuation of the response over time rather than a narrowing of the response bandwidth. (**C**) Amplitude of band-limited activity in three frequency bands for the same electrode. Shading corresponds to standard error. While including lower frequencies increases the response SNR, all subsets of the high-frequency range produce a qualitatively similar response pattern, suggesting that the underlying activity is indeed broadband.

### Duration-Tracking Decreases from EVC to Category-Selective IT Cortex

To determine the degree to which the real-time, moment- by-moment presence of the visual percept is analogically reflected in the HFB signal on a single-trial basis, we quantified for each trial and electrode the duration for which the temporally-smoothed HFB signal was continuously sustained above baseline following the response onset. A *duration-tracking accuracy* was then computed as the percentage of trials where the duration of the sustained response was within ±150 ms of the true stimulus duration (see Materials and Methods). The rationale of this approach is to emulate a real-time process where the persistence of the stimulus percept can be decoded from the region’s HFB signal, by testing, at each consecutive time window, whether the signal is still above the decision threshold (“activation”) or not (“baseline”). Of the 292 visually-responsive electrodes, 21 electrodes in eight subjects showed significant duration-tracking accuracy based on a permutation test, 17 of which were located in the posterior occipital lobe (50% of responsive early visual cortex electrodes) and 4 located on IT cortex (1.5% of responsive IT electrodes; Fig. 3A, 5).

**Figure 3:**
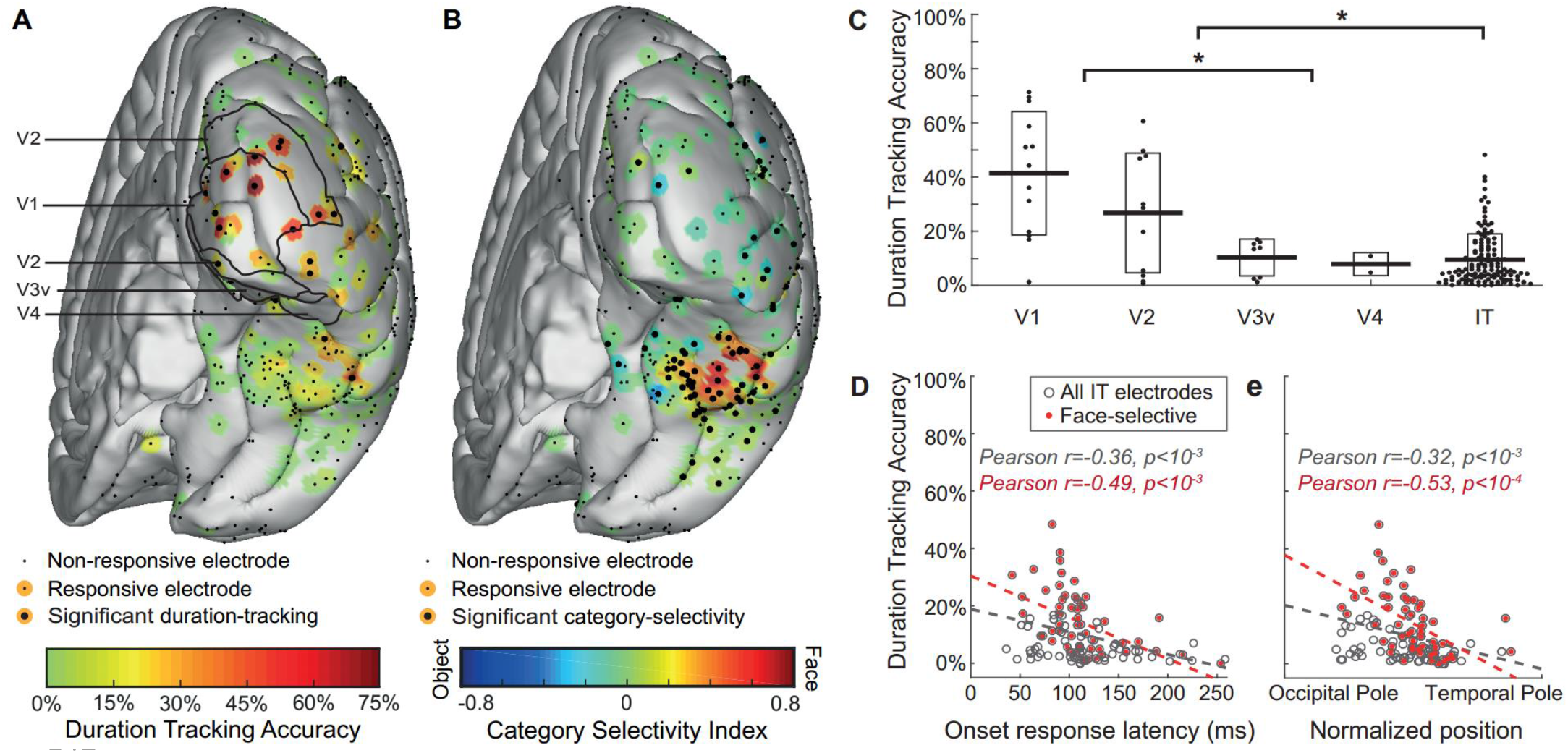
Duration-tracking accuracy decreases gradually along the ventral visual stream. (**A**) Duration-tracking accuracy for all subjects’ electrodes projected onto a common brain template. Visually responsive electrodes are surrounded by a color patch representing duration-tracking accuracy. Significant duration-tracking (FDR corrected, q = 0.05) is denoted by a thicker black dot. Anatomical regions marked with black contour lines within EVC are based on surface registration to a probabilistic atlas (Wang et al. 2014). (**B**) Same as (A), but for category selectivity index. Negative values (blue) correspond to object-preference. (**C**) Relation between duration-tracking and hierarchical position along the ventral stream (EVC areas defined based on the probabilistic atlas). Each dot corresponds to a single electrode, with horizontal dispersion based on data point density. Boxes correspond to standard deviation. Asterisks mark significant difference (p<0.05) between EVC areas and IT and between V1/V2 and V3v/V4. Note that V3v/V4 sites are not necessarily earlier than all IT sites in terms of response latency. (**D**) duration-tracking within IT plotted against onset response latency, as a proxy for hierarchical position along the ventral stream. Face selective electrodes are marked red. (**E**) Same as (D), with hierarchical position measured as the electrode’s coordinate along the occipital-temporal axis.

To identify category-selective electrodes, a selectivity index (SI) metric was computed corresponding to the response amplitude difference between faces and objects in the 300 ms post-onset window, normalized by their sum (positive SI corresponds to face-preference; see Materials and Methods). One hundred and twenty seven electrodes across all subjects showed category-selectivity, 96 of which were on IT (64% of IT electrodes), and the rest on the occipital, parietal, or inferior frontal lobes.(Fig. 3B). In six category-selective electrodes that were also duration-tracking, we additionally tested whether category selectivity was sustained beyond the onset response, by calculating the SI in the 1200-1500 ms post-stimulus window for 1500-ms-duration stimuli. Significant selectivity in sustained activity was found in two duration-tracking electrodes in posterior IT.

As expected, category selectivity was low over EVC and maximal over the fusiform gyrus. In contrast, duration-tracking accuracy was maximal over EVC and followed a diminishing gradient along the ventral stream toward anterior IT – such that the majority of responses in regions highly informative about the category of a stimulus showed low sensitivity to stimulus duration, and thus to its presence beyond its onset (Fig. 3).

EVC electrodes showed significantly higher duration-tracking accuracy than IT electrodes (Fig. 3C, t(166)=7.01, p<10^−5^). Within EVC, V1-V2 electrodes showed higher accuracy than downstream V3v-V4 (t(32)=3.31, p<10^−3^1). There was a trend for higher accuracy in V1 than in V2 (t(22)=1.59, p=0.06). Within IT, we tested for a posterior-anterior gradient using onset response latency and anatomical location as two converging indexes for position along the ventral stream. Anatomical position was quantified by projecting the coordinates of each electrode to a line extending from the occipital pole to the temporal pole of each individual subject’s right hemisphere, and normalizing the result to the range 0-1 (1 is closer to the temporal pole; electrodes over the left hemisphere were omitted from this analysis due to their paucity). Duration-tracking accuracy was negatively correlated with response latency (Fig. 3D, r=-0.36, p<10^−3^) as well as position (Fig. 3E, r=-0.32, p<10^−3^). The same correlations were found when limiting the analysis to face-selective electrodes (r=0-.49, p<10^−3^ for onset latency and r=-0.53, p<10^−4^ for position). Object-selective electrodes showed a similar negative correlation but were too few for robust statistical results. The reported results were unchanged when controlling for signal-to-noise ratio across electrodes as expressed by onset response peak magnitude and baseline noise level (Fig. S1), or when using a mixed-model regression approach to account for differences between subjects (Fig. S2). Similar results were obtained when the mean error of the response estimation (i.e. difference from true stimulus durations) was used as dependent variable instead of the duration-tracking accuracy metric (Fig. S3), or with a more liberal index of *duration-dependence*, based on the correlation between the stimulus duration and the number of above-threshold post-stimulus time points across trials (Fig. S4). Sites with low duration-tracking accuracy did not always show only transient onset responses. Rather, many non-duration-tracking electrodes were characterized by prolonged HFB activity that was uninformative about single-trial stimulus duration. Fig. 4 shows representative examples of an EVC electrode with high duration-tracking accuracy (left), a face-selective IT electrode with a prolonged response that is not informative regarding the duration of the stimulus (right), and a rare posterior IT electrode with a category-selective, duration-tracking response (middle). We next describe this rare response profile, encoding both the content and the ongoing presence of a stimulus over time.

**Figure 4:**
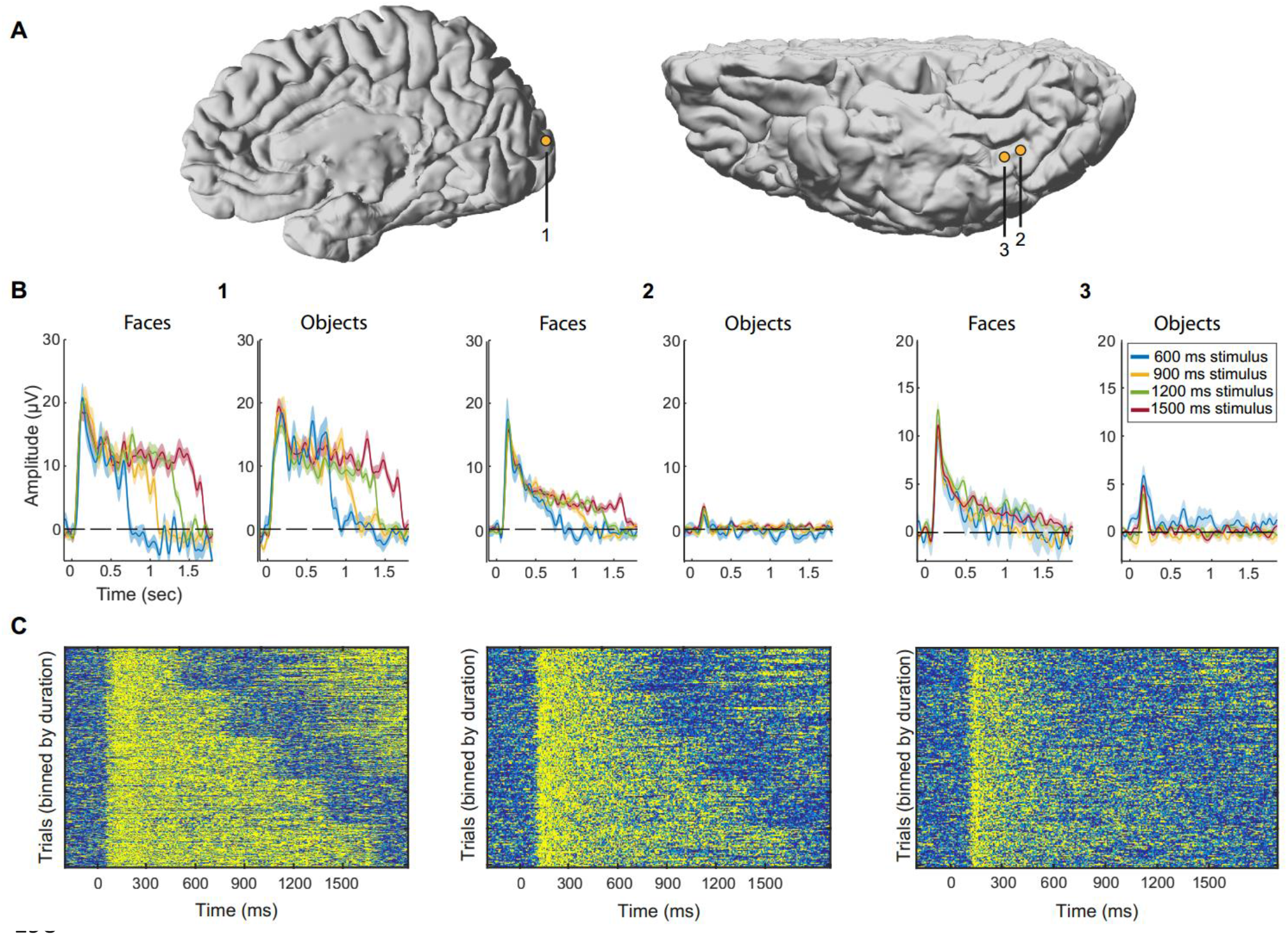
Three representative electrodes. Electrode 1 – Early visual cortex electrode (dorsal V2/V3 according to fMRI retinotopy performed on this subject for another study, see Parvizi et al. 2012), duration-tracking but not category-selective (trials pooled from both categories). Electrode 2 – Lateral fusiform gyrus electrode with sustained activity that is both duration-tracking and category-selective (face trials only). Electrode 3 – Slightly anterior to electrode 2 on the lateral fusiform gyrus, category-selective but not duration-tracking. MNI coordinates for the three electrode are (9.4, −94.8, 12.7), (37.6, −29.5, −11.1), (39.1, −25, −11.2). (**A**) Electrode locations on native brains (electrodes 2, 3 are from the same subject). (**B**) Mean response for each stimulus duration and category. For clarity only trials lasting at least 1800 ms (onset to onset) are shown, thereby eliminating 300 ms stimulus duration trials. Shading corresponds to standard error across trials. Note different y-axis scales. (**C**) Single-trial images (stacks) binned by stimulus duration (300, 600, 900, 1200, 1500 ms from top to bottom). Each row in the images represents a single trial. Increased activation at the end of short-duration trials corresponds to the subsequent trial when the ISI is short.

### Stimulus Presence and Category are Jointly Encoded in Posterior IT Cortex

Two face-selective electrodes from two subjects showed both duration-tracking and sustained category selectivity (Fig. 5). Both were similarly located laterally from the posterior part of the mid-fusiform sulcus (Weiner et al. 2014). Two additional posterior fusiform electrodes, located medial to the mid-fusiform sulcus, were also object-selective and duration-tracking, however their selectivity was limited to the onset response, that is, the response to faces and objects did not differ in the late (1200-1500 ms) time window. The locations of the two face-selective electrodes correspond to the posterior fusiform face-selective region pFus-faces (or FFA-1, Grill-Spector & Weiner 2014), and to cytoarchitectonic region FG2 (Caspers et al. 2013), suggesting a functional distinction between this region and the more anterior mFus-faces region (FFA-2, related to cytoarchitectonic area FG4, Lorenz et al. 2015), which showed high category selectivity but little duration-tracking.

### Differences In Duration-Tracking Are Not Explained By Task Attentional Demands

The diminished sustained activity in IT could be explained by the possibility that subjects stopped attending to the stimulus or averted their gaze after its appearance (eye tracking was not feasible at the ICU bedside). If IT is more sensitive than EVC to the level of attention, this could drive the dissociation between these regions in sustained responses (Levy et al. 2001; Hasson et al. 2002). To test this, a second version of the experiment was performed by 7 of the 10 subjects in addition to the original version. In this difficult dual-task control, they had to detect a rare brief blurring of any image just before its disappearance, in addition to the detection of clothing items (Fig. 1B). The dual detection task required gaze and attention to be sustained throughout image presentation as the timing of stimulus offset was unpredictable. Detecting clothing items in the dual-task control experiment provided similar results to the main single-task experiment for all subjects. Two subjects with relevant category selective IT electrodes (8 and 21 electrodes over the left IT cortex) adequately performed the second, blurring detection task in the dual-task control experiment (hit rates 75% and 94%; hit rates for detection of clothing items in these subjects: 88% and 94%; overall false alarm rates 4% and 1% respectively). The same relation between HFB and non-target stimulus duration was found with the dual-task control as with the single-task original experiment in the IT electrodes from these two subjects, and well as in all other electrodes for all seven subjects (Fig. 6; the other subjects either missed the majority of the blurring targets while continuing to respond to the clothing items, or did not have suitable electrode coverage for this analysis). This argues against the possibility that the absence of sustained activity in IT electrodes was due to lack of sustained spatial attention, or shifts of gaze away from the stimulus. Indeed, while it could be expected that the introduction of the secondary control task will increase activity along the visual pathway because of the higher attentional demands (Brefczynski & DeYoe 1999), subjects likely paid sufficient attention to the stimuli already in the main experiment (creating a ceiling effect for attention). Given this insensitivity to experimental condition, data from both experiments was pooled in all other analyses.

**Figure 5:**
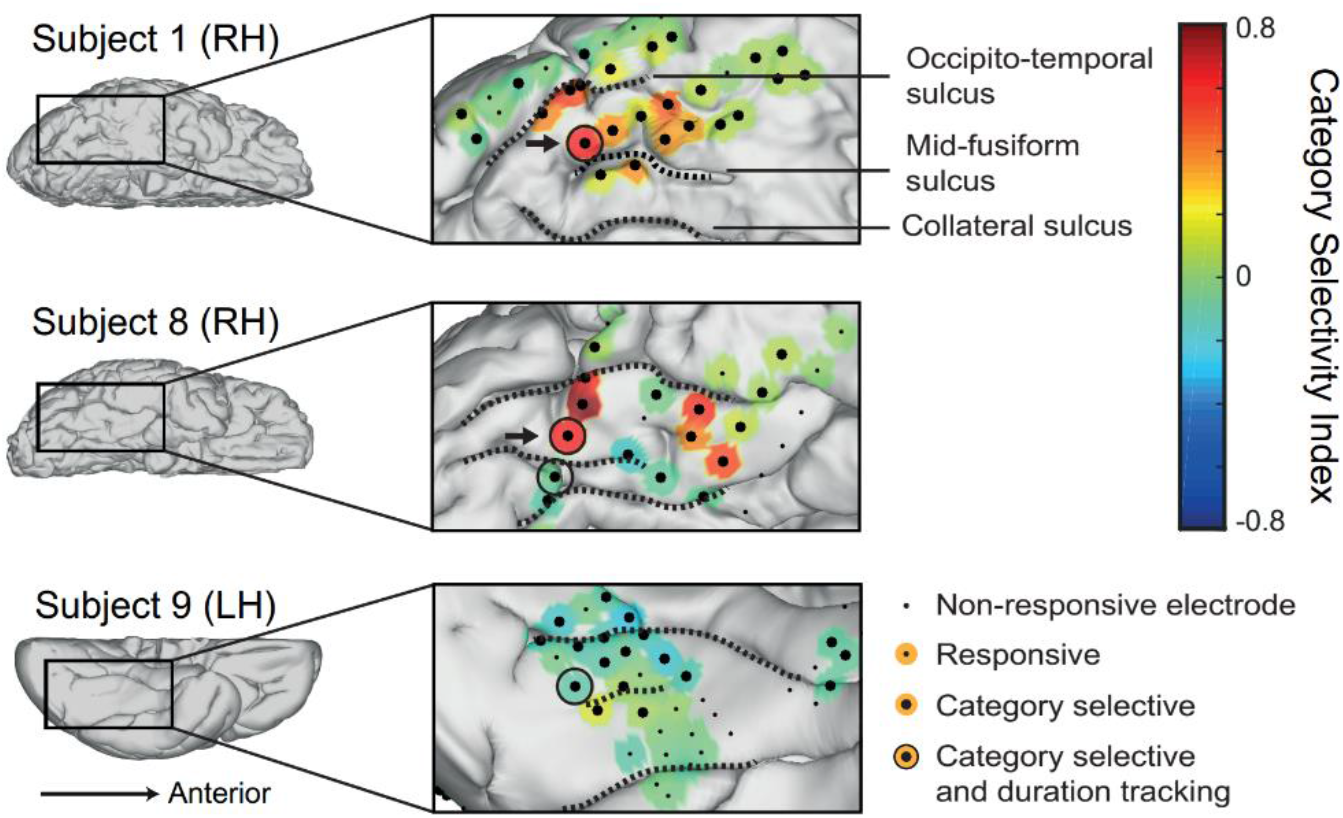
Electrodes on the posterior fusiform gyrus encode stimulus persistent presence and its category. Four significantly duration-tracking electrodes in three subjects were found within category-selective IT cortex, all located adjacently to the mid-fusiform sulcus on the posterior fusiform gyrus, as opposed to more anterior category-selective sites which do not track ongoing stimulus presence. Of these, the two face-selective electrodes (from two subjects) marked by arrows also maintained category-selectivity persistently throughout the duration of the response (marked with arrows) whereas the two others did not. The electrodes are presented on the subjects’ native brains.

**Figure 6:**
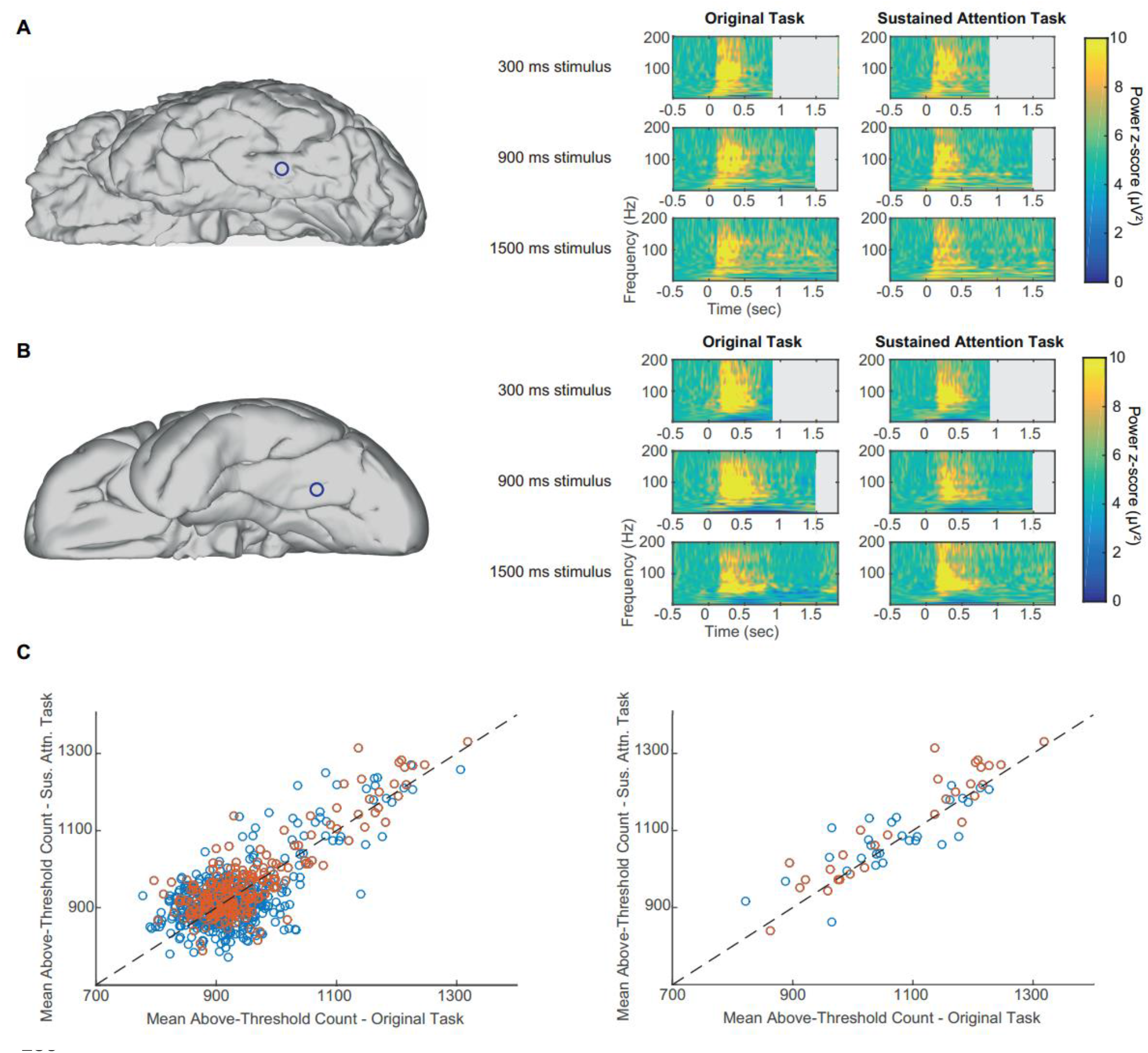
No differences between single task and dual-task attention control. Panels A,B show time-frequency plots from two representative electrodes in two subjects who performed well for both the clothing items and the blurring events detection task (attention-control). Left column: electrode location on individual subject’s brain. Right column: time-frequency plots for a single electrode corresponding to three stimulus durations (top to bottom) and the two experimental conditions. Power is shown as z-score compared to baseline. (**A**) Face-selective, non-duration-tracking IT electrode. (**B**) Object-selective, non-duration-tracking IT electrode. (**C**) Left: each electrode is plotted as one data point comparing the mean number of above-threshold data points (across trials) between the two tasks (r=0.74). The results are clustered around the diagonal demonstrating similar mean response durations across conditions. Orange points are from the three subjects who could perform the control task well. Right: the same relation is shown specifically for category-selective inferior temporal electrodes. There was no significant difference between the two experiments either on the single electrode basis (across trials) or in the mean across electrodes.

### Duration-Tracking Activity Is Not Driven By Eye-Movements

The finding of duration-tracking responses could be potentially explained as the summed activity of transient (onset) responses to retinal shifts driven by saccadic eye movements (EM) while the stimulus is presented (Nagasawa et al. 2011). However, there are good reasons to suggest that duration-tracking responses are not merely driven by EM. First, eye movements occur about 1-2 (max. 4) per second, and thus EM-driven activity would have taken the form of random sparse bursts in single trials, but these were not consistently observed. Second, although eye movements could not be tracked with fidelity in the clinical setting at the time the data were collected, saccadic spike potentials elicited by the contraction of the ocular muscles at the onset of saccades (Yuval-Greenberg et al. 2008), can be detected intracranially in electrodes proximal to the eyes (Kovach et al. 2011; Jerbi et al. 2009). We were able to identify reliable saccadic events in one of our patients through the detection of intracranial saccadic spike potentials in two temporal pole electrodes, just posterior to the eye (Fig. 7A). We detected saccadic potentials using a custom filter, based on a canonical spike potential template derived from independent recordings of saccadic spike potentials from three EEG control subjects who performed the same experiment with concurrent eye-tracking (see Materials and Methods). The claim that the intracranial spikes indeed corresponded to saccadic potentials was supported by producing a peri-stimulus saccade rate modulation curve, which matched the pattern observed in the eye-tracked control subjects and was consistent with expected peri-stimulus pattern for saccades (i.e. initial suppression followed by increased saccade rate, Fig. 7C), as well as by the theoretically expected mean saccade rate of about 1.5 saccades/sec (Engbert 2006). Finally, to test for the effect of saccades on sustained high-frequency visual responses, we examined data from a posterior face-selective duration-tracking inferior-temporal electrode, in the same subject in which the intracranial spike potentials were recorded. The data was divided into two trial groups: one group of 11 trials having zero detected saccades within the 1500 ms post-stimulus window, and a second group of the same number of trials having the most detected saccades (3-6 per second, mean 4.2). The HFB response to 1500-ms-duration face stimuli was then averaged for each group and compared. The results show no difference between the trial groups (Fig. 7D,E). Similar results were obtained by comparing the trials with more saccades to those with less saccades by a median split based on the number of saccades per trial. Thus, face-selective duration-tracking in a posterior IT electrode was not contingent on the presence of saccades during stimulus presentation. Finally, to further test whether the sustained activity in early visual areas could be explained by saccade-locked activity, we simulated a best-case scenario for this hypothesis and tested it against the empirical results (Fig. S5). The simulation results show that saccade-driven responses could not account for the robust duration-tracking activity observed in early visual cortex.

**Figure 7:**
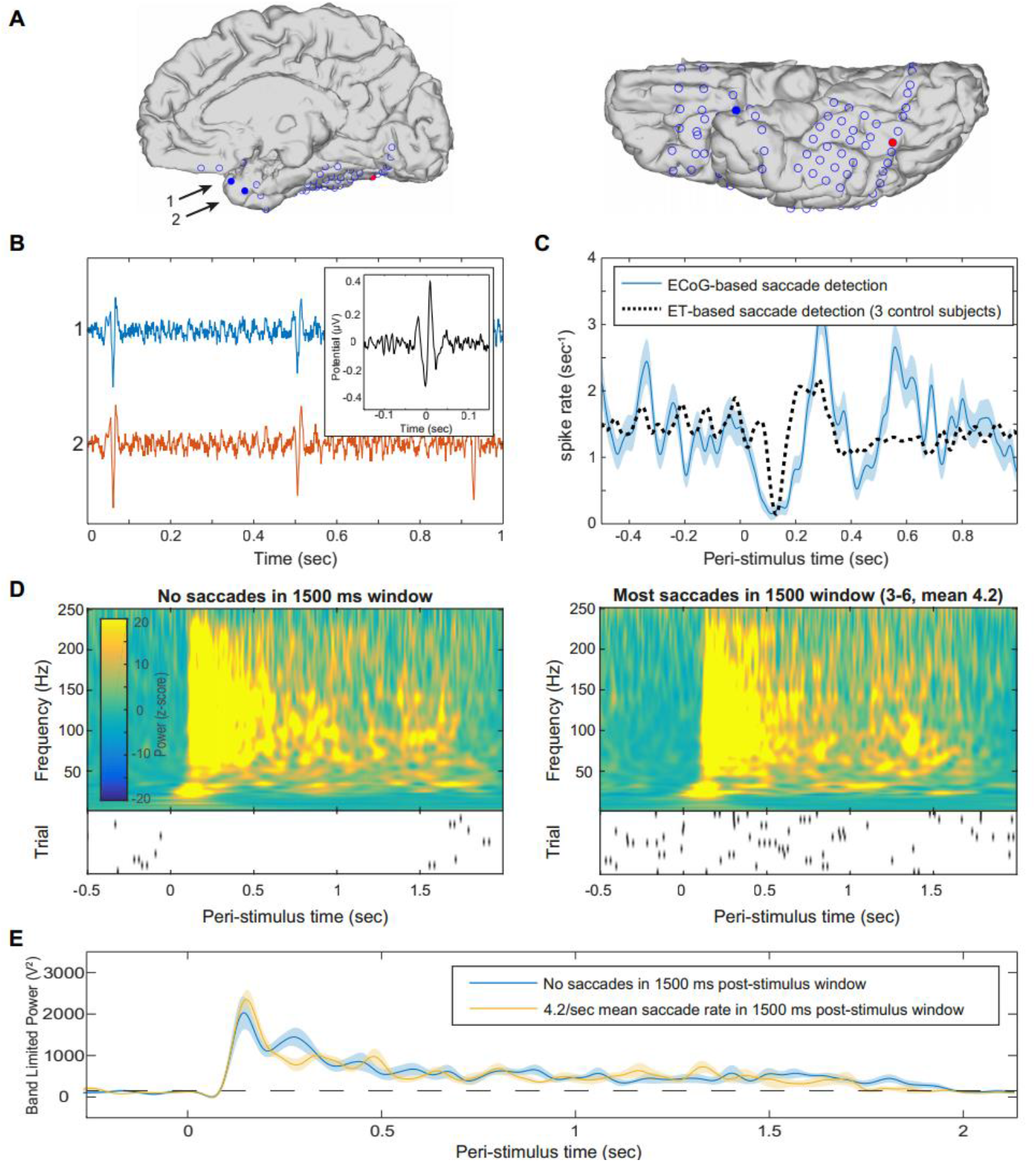
No effect of saccades on sustained visual activity. Saccadic events were identified in a single subject based on detection of saccadic spike potentials in two electrodes near the temporal pole. (**A**) Anatomical location of electrodes used to detect saccadic spikes (blue filled circles), and an electrode in the same subject exhibiting face-selective duration-tracking (red). (**B**) Representative un-averaged traces (>30 Hz high-pass-filtered) of spike events in the two temporal-pole electrodes. Inset: the averaged saccadic spike model from the 3 EEG control subjects which served as a model and in which saccade onsets were detected by eye tracking. (**C**) Peri-stimulus saccade rate. The ECoG-based saccades follow a typical post-stimulus suppression and rebound (blue), comparable to saccades detected using eye-tracking for the same task in control subjects (black dotted). (**D**) Average time-frequency plots of the responses to 1500 ms face stimuli in the posterior IT electrode showing face-selective duration-tracking, for trials with no saccades during stimulus presentation (left), vs. trials with many saccades (right). Shown below each time frequency plot is a raster plot of saccade events during each trial included in the group. Each row represents a single trial and each tick mark represents a detected saccade. (**E**) Direct comparison of high-frequency band-limited power for the two groups of trials. Cluster-based permutation testing did not detect differences in power at any time point. Dotted line marks mean baseline power.

### Additional Manifestations of Duration Dependence

In addition to the analysis of the HFB signal, duration-dependence was also assessed in the time domain (i.e., event related potential) using the same method as for HFB, except that original rather than bootstrapped trials were used and the signal was inverted in polarity if the mean of the 1000-1500 ms window of the response to 1500 ms duration stimuli was negative, to allow for negative sustained potentials. We found positive and negative *duration dependent sustained potentials* (DDSPs), that is, deflections which remained distinct from baseline for the duration of the stimulus (cf. N’Diaye et al. 2004 in a duration-estimation task). These were found in four non-category-selective electrodes on the inferior bank of the calcarine sulcus in three subjects (Fig. 8), all mapped to ventral V1 based on a probabilistic atlas (Wang et al. 2014). Of the four electrodes with significant DDSPs, three showed also significant HFB duration-tracking and one only a non-significant but visually discernible sustained HFB response. In contrast, other sites with strong HFB duration-tracking in the same subjects did not show any type of DDSP. We note that in some of the subjects the 0.5Hz high pass filter applied during the recording might have reduced the sensitivity to DDSPs.

**Figure 8:**
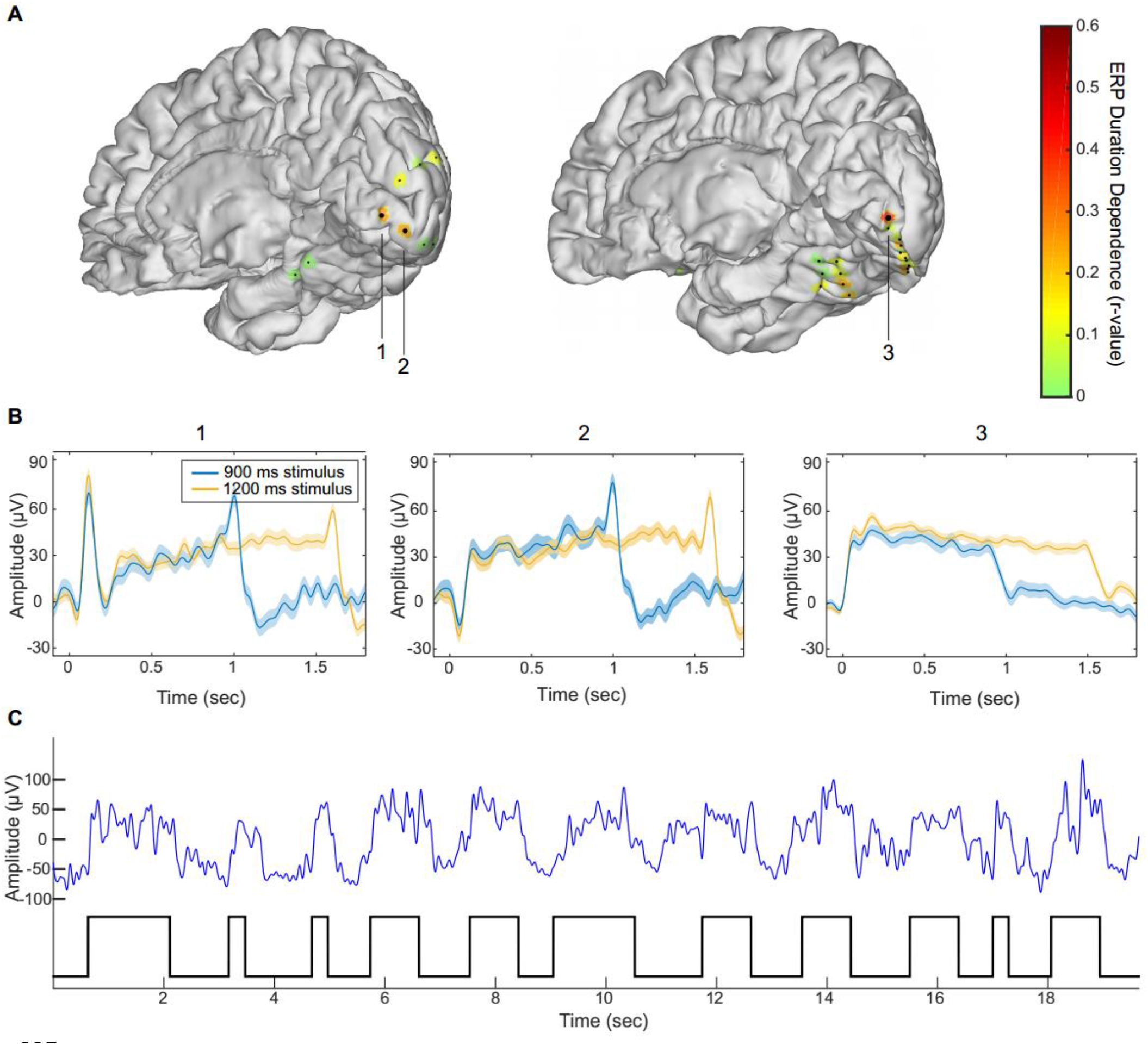
Duration dependent sustained potentials. (**A**) Duration-dependence of evoked responses in two individual subject brains (subjects 5 and 10, middle and right). Colored patches indicate degree of duration-dependence according to the color scale on the right. Large dots indicate statistically significant duration-dependence. (**B**) Mean evoked responses for the three electrodes identified in panel (A). (**C**) High-SNR segment of the raw data trace (blue line, low-pass filtered at 10Hz) taken from electrode 3 and on-off visual stimulation epochs (black trace on bottom). The robust DDSP evidenced without averaging, indicates that DDSP corresponds to sustained single trials deflections rather than to a summation of stimulus-contingent transient deflections in single trials.

Additional analyses of cross-frequency coupling and inter-electrode phase-coherence did not uncover any duration-dependent behavior (see Materials and Methods). Duration-dependence could also characterize the power modulation of low-frequency bands (e.g. in the alpha/beta range). However, because of the strong low-frequency power of the onset and offset responses, and the lower temporal resolution of time-frequency analysis in these bands (temporal smearing), resolving low frequency duration-tracking necessitates longer stimulation durations than were used in our study.

## Discussion

### Duration-Tracking as an Organizing Principle in the Ventral Stream

We found that while activity in early visual cortex remained above threshold for as long as the stimulus was presented, this correspondence diminished along the visual hierarchy and was minimal in category-selective sites in anterior IT cortex. This anterior-posterior gradient was evident through a direct comparison of duration-tracking accuracy between anatomically-defined visual regions, as well as its correlation with onset response latency and with electrode coordinates on a posterior-anterior vector.

The functional organization of visual IT cortex obeys a number of well-characterized functional-anatomical regularities. These include an increasingly abstract, high-level representation (Lerner et al. 2001; Kanwisher 2010), increasing spatial receptive field (Wilson & Wilkinson 2015) and temporal (Honey et al. 2012) receptive window size along the posterior-anterior axis, as well as variable preference for stimulus eccentricity (Hasson et al. 2002), animacy, and stimulus size (Weiner et al. 2014) along the medial-lateral axis. Our results add temporal dynamics as another organizing principle, manifested as diminishing real-time perceptual duration tracking along the visual hierarchy. This gradient is comparable in nature to the other posterior-anterior gradients in that it entails a growing invariance to low-level properties of the visual signal.

From a theoretical perspective, this gradual attenuation of information about stimulus presence may reflect a basic property of information transfer along a serial processing stream. As formally stated by the data processing inequality theorem (Cover & Thomas 1991), in any system the amount of information about the initial input cannot be increased as a result of additional processing. Thus, any information about the real-time presence of a visual percept in a cortical site must be present also in the input to that site. Given imperfect synaptic transmission, this information can be expected to become less reliable along the processing stream – and likely more so for sustained than onset-driven activity, as the former is typically of lesser magnitude making it more likely to fall below neural detection thresholds^*^. Persistent stimulus-driven neural activity corresponding to the presence of a visual feature at any processing stage can thus be expected to be only as reliable as the persistence of its input from the previous stage, and as such is bound to show some degree of decay with subsequent processing steps. Functional differentiation in visual processing along the ventral stream could in part arise from this type of fundamental property of the cortical system.

### Differentiation Within Category-Selective Cortex

We found that within face-selective regions of the fusiform gyrus, significant duration-tracking activity was found only at some posterior recording sites, corresponding to the FFA-1 region, while more anterior (FFA-2) sites showed non-reliable duration-tracking activity. These results provide evidence that regions which appear homogenous in terms of their category-selectivity (Tsao et al. 2006), but are anatomically and cytoarchitectonically distinct (Caspers et al. 2013; Lorenz et al. 2015; Grill-Spector & Weiner 2014), differ in how they respond to a stimulus over time and in their potential contribution to ongoing visual perception. This differentiation is of particular interest since the temporal response profile of a cortical site presents a constraint on its functional role. For example, cortical activity showing reliable (single-trial) duration-tracking may maintain an ongoing visual percept, while transient responses can be a signature of stimulus-onset-related processes such as categorization or identification.

The temporal response profile difference found between EVC and IT regions and between posterior and anterior category selective regions could be related to previous fMRI results (Weiner et al. 2010), although the paradigm and especially the time scales are quite different. In Weiner et al.'s study, IT regions showed larger fMRI adaptation effects (reduced BOLD response to repeated vs. non-repeated stimuli) than EVC regions, and anterior face- and limb-selective regions showed stronger adaptation than their more posterior counterparts. The latter difference was pronounced in a condition with long lags between repetitions (many seconds and intervening stimuli) and not as much with short lags (half a second). The present study, using direct neural measures and high temporal resolution, may have been more sensitive to differences between regions at the sub-second time scale. Moreover, it suggests that a gradient of temporal response profiles is present even within the response to a single stimulus, and not only in the adaptation across stimuli.

### Measuring Neural Activity Under Sustained Cognitive Conditions

The challenge of measuring neural activity contingent on a sustained cognitive process, while dissociating it from other temporal activity profiles such as transient onset- or offset-driven activity, has not been widely addressed in the literature. A common approach in functional imaging studies of sustained cognitive processes (e.g. working memory, sustained attention or imagery) is to use separate GLM predictors for onset-, sustained- and offset-/response-related activity (Eriksson et al. 2004; Portas et al. 2000; Curtis & D’Esposito 2003; Offen et al. 2009). However, this does not produce reliable evidence for sustained activity that is independent from onset and offset responses, because any neural activity extending beyond the duration of the modeled onset response will be interpreted as sustained activity, regardless of whether its duration is in fact related to the duration of the process. The current study suggests two general guidelines for measuring duration dependent sustained activity: first, the experimental paradigm should allow for variable response durations; and second, the analysis should directly assess the dependence of the response duration on the experimental duration (see Vilberg & Rugg 2012 for an example of such analysis in the context of memory retention). Direct examination of duration-tracking in neural activity associated with sustained cognitive processes may benefit a wide range of studies.

## Summary

Onset responses reflect major changes in the environment, yet much of perception occurs between changes. We addressed the neural basis of sustained visual perception and report dissociations between semantic category ("What") and ongoing presence ("Whether”) of a visual event evident along the ventral visual pathway. Understanding the neural correlates of perception beyond its onset has major implications for deciphering the biological underpinning of experience.

## Materials and Methods

### Subjects

Ten patients (3 female, 7 male, mean age 41.5 years, range 19-65, two left-handed) undergoing surgical treatment for intractable epilepsy and implanted with chronic subdural electrodes participated in the experiment. Seven patients were recorded in Stanford School of Medicine, 2 in the California Pacific Medical Center (CPMC), and 1 in the UCSF Medical Center. Electrode arrays were implanted on the right hemisphere for 8 of the patients, and on the left hemisphere for 2, with electrode location determined solely by clinical needs. All subjects gave informed consent approved by the UC Berkeley Committee on Human Research and corresponding IRBs at the clinical recording sites.

### Stimuli and Tasks

Recording was conducted in the epilepsy ICU. Stimuli were presented on a laptop screen and responses captured on the laptop keyboard. Stimuli were grayscale images of either frontal human faces, round watch-faces, other man-made objects (tools, vehicles, furniture, musical instruments and household objects), or animals, presented on a square uniform gray background in the center of the screen and extending across approximately 5° of the visual field in each direction. For the analyses reported here, watch-faces and other man-made objects were grouped together as “objects”, and only the face and object categories were used in the main analyses. Analysis for the animal category is reported in Fig. S6. Additional categories including body parts and houses were excluded from analysis due to consisting of too few exemplars. Face and object images did not significantly differ in terms of luminance (206.2 and 207 mean pixel intensity for faces and objects, respectively) or contrast defined as the standard deviation of pixel intensity (41.8 and 41.1 respectively). Each stimulus was presented for either 300, 600, 900, 1200 or 1500 milliseconds for the first 3 subjects, or 300, 900 or 1500 milliseconds for the last 7 subjects (the number of durations was reduced to allow the additional control task described below without excessively prolonging the experiment). All categories had the same probability of appearing with each duration. Inter-stimulus interval varied from 600 to 1200 ms with 150 ms steps, during which a fixation cross was presented. In the main experimental condition, subjects were instructed to fixate on the center of the screen, and respond with a button press whenever a clothing item was presented (10% of trials). The last 7 subjects performed an additional experimental task (dual attention-control task) where in addition to responding to images of clothing items, they were also required to respond to rare instances when an image of any category became blurry for the last 200 ms of its presentation (clothing targets and blurred image targets each accounted for 5% of the trials). The onset time for each image was registered alongside the ECOG data from a photodiode placed on the laptop screen, recording a white rectangle displayed at the same time as the image at the corner of the screen.

### ECoG Acquisition and Data Processing

Each subject was implanted with subdural arrays containing 53-128 contact electrodes (AdTech Inc.). In total, 1067 electrodes were examined. Each electrode was 2.3 mm in diameter, with 5 or 10 mm spacing between electrodes within an array, arranged in 1-dimensional strips or 2-dimentional grids. Recordings were sampled at 1000 Hz (CPMC), 3051.76 Hz (Stanford, UCSF) or 1535.88 Hz (Stanford) and resampled to 1000 Hz offline (with the exception that the original 3051.76 Hz sampling rate was used for the results reported in Fig. 2). A high-pass filter was applied online to the signal at either 0.1Hz or 0.5Hz in different subjects. 159 electrodes manifesting ictal spikes or persistent noise were visually identified and removed from analysis, as were time intervals with excessive noise or ictal activity as determined by one of the authors (RTK). All remaining electrodes were re-referenced offline to the average potential of all non-rejected electrodes, separately for each subject. Unless otherwise indicated, all data processing and analysis were done using custom Matlab code (Mathworks, Natick, MA).

### Electrode Localization

Electrode locations were identified manually using BioImageSuite (www.bioimagesuite.org) on a post-operative Computed Tomography (CT) scan co-registered to a pre-operative MR scan using the FSL software package (Jenkinson & Smith 2001; Jenkinson et al. 2002). Individual subjects’ brain images were skull-stripped and segmented using FreeSurfer (http://surfer.nmr.mgh.harvard.edu). Localization errors driven by both co-registration error and anatomical mismatch between pre- and post-operative images were reduced using a custom procedure which uses a gradient descent algorithm to jointly minimize the squared distance between all electrodes within a single electrode array/strip and the cortical pial surface (see Dykstra et al. 2012 for a similar procedure). In contrast to methods which only attempt to correct individual electrodes’ position in relation to the pial surface, this method preserves the original array topography, thus providing a more reliable estimate of actual electrode positions.

Individual patients’ brains and electrode coordinates were next co-registered to a common brain template (FreeSurfer’s *fsaverage* template) using surface-based registration (Fischl et al. 1999). This registration optimally preserves the mapping of electrode locations to anatomical features between the native and common brain space despite individual variability in cortical folding patterns, in contrast to volume-based registration which preserves the 3D spatial configuration of electrode arrays but does not account for differences in cortical folding patterns. To identify visual areas, each cortical surface was resampled to a standard mesh with a fixed number of co-registered nodes using SUMA (Argall et al. 2006), and visual area labels were imported from a probabilistic map of visual areas topography (Wang et al. 2014). Visualization of electrode data in Fig. 3 was based on surface registration to an MNI152 standard-space T1-weighted average structural template image.

### High Frequency Broadband Power Signal

To extract the high-frequency broadband (HFB) signal, the recorded signal was high-pass filtered using a 3^rd^ order Butterworth filter with a 30Hz cutoff, and the power along time in each electrode was computed using the square of the Hilbert transform of the filtered signal. The >30Hz frequency band was initially selected for further analysis based on visual inspection of the spectral responses for all stimuli at each electrode, averaged across all categories and durations. Morlet wavelet analysis was used solely for visualization purposes (Fig. 2, 6, 7), with a wavelet constant of 12 and center frequency bins ranging from 2 to 800Hz.

### Identification of Visually-Responsive Electrodes

Visually-responsive electrodes were defined as those exhibiting a significant (p<0.01) HFB power increase in response to either the face or object stimulus categories across trials, as assessed by a cluster-based permutation test compared to a 500 ms pre-stimulus baseline (Maris & Oostenveld 2007). Subsequent analyses were performed on the set of electrodes showing a significant response to at least one category. In addition, the response onset latency of each visually responsive electrode was derived as the onset latency of its earliest significant activity cluster.

### Category-Selectivity

Category selectivity was defined as a significant difference between the HFB onset response to face vs. object stimuli (note that this type of selectivity does not necessarily imply specificity to the preferred category, and could arise out of a less specific selectivity, e.g. animate vs. non-animate stimuli (Grill-Spector & Weiner 2014) – for an analysis aimed at achieving greater category specificity based on comparison to a third category see Fig. S6). The onset response magnitude for each category was computed by taking the root mean square (RMS) of all data points from the first 300 milliseconds of the trial-average HFB response to the category. A selectivity index (SI) was computed as:

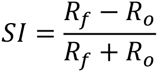

Where *R_f_* and *R*_*0*_ are the RMS of the mean response to faces and objects, respectively. Positive values correspond therefore to face-preference, and negative values to object-preference. Statistical significance was derived with a randomization test whereby in each of 10,000 iterations the category labels were shuffled and the SI recomputed, producing a null-hypothesis distribution. P-values were derived from the rank of the original SI within this distribution (in absolute value), corrected for multiple comparisons across all electrodes using the False Detection Rate (FDR) method (Benjamini & Yosef 1995). In this analysis and whenever FDR was used in this study, the FDR q value was 0.05.

### Single-Trial Duration Tracking

The purpose of the single-trial duration tracking analysis was to identify electrodes with singletrial HFB responses which are reliably (across trials) sustained above baseline for a duration corresponding to the stimulus duration (i.e. duration-tracking responses). The rationale of this approach is to emulate an online process where the persistence of the stimulus percept can be decoded from the region’s current HFB signal, by testing at each consecutive time point whether the signal is still above the decision threshold (“activation”) or not (“baseline”).

The HFB signal was first smoothed using a 300 ms window moving average. To derive the activation threshold, an “activation” sample was generated by taking the means of the 1200-1500 ms post-stimulus window across 1500-ms-duration trials, and a “baseline” sample was generated by taking the means of the 300 ms pre-stimulus window from the same trials. The late 1200-1500 ms window was selected to allow the test to be sensitive to the relatively low-magnitude activation of the late sustained response as opposed to the high-magnitude early response component. The threshold was then set as the median value of the two equally-sized samples taken together. For each trial, the threshold was generated based on all trials excluding the current one. In summary, the threshold indicates whether a power value averaged across 300 ms is more likely to reflect a baseline or active epoch. The decoded duration of the active percept was then defined as the latency (relative to the electrode’s onset response latency) of the first time point where the smoothed HFB signal drops below the threshold, within a 1800 ms post-stimulus period. Only trials which included a minimal 1800 ms window from stimulus onset to the next trial stimulus onset were included in this analysis, thus excluding all 300 ms duration trials, and some of the 600 and 900 ms duration trials. Using a shorter analysis window of 1200 ms and thus including all 300, 600 and 900 ms trials, while excluding 1200 and 1500 ms trials, led to very similar results. Finally, the duration-tracking accuracy was computed as the percentage of “correct” estimated durations which were within a ±150 ms error margin around the true stimulus duration. The results of this analysis were not qualitatively changed when selecting a different error margin (i.e. ±100 or ±200 ms), or smoothing window size (i.e. 200 or 400 ms). Statistical significance was determined using a randomization test such that at each of 10,000 iterations, the duration-tracking accuracy was re-computed based on shuffled stimulus duration labels to produce a null-hypothesis distribution. P-values were derived from the rank of the original accuracy within this distribution, corrected for multiple comparisons using the FDR method. Duration-tracking was assessed across both face and object stimulus trials for non-category-selective electrodes, and for the preferred category for category-selective electrodes.

### Duration-Dependence

Duration-tracking accuracy is a strict index of sustained activity as it requires that HFB does not drop below baseline throughout the stimulus duration in single trials. We also derived a more sensitive index corresponding to a weaker notion of *duration dependence* by correlating trials’ duration with the number of above-threshold HFB data points in a post-stimulus window. This resulted in a correlation coefficient for each electrode, where 0 indicates no statistical relation between stimulus duration and the number of above-threshold HFB time points and 1 indicates a perfect linear relation. To determine the number of above-threshold points, first the HFB signal was smoothed using a 3^rd^ order Butterworth 10 Hz low-pass filter, and trials were baseline-corrected by subtracting the average of the 500 ms pre-stimulus period. To maximize the sensitivity of this metric, a bootstrapping method was applied whereby for each duration, 1000 surrogate trials were generated by each time averaging 10 randomly-drawn trials of the same duration. This averaging ensured that any real difference in the response to different durations would not be concealed by single-trial noise. Next, an “activation threshold” for each electrode was set as the median of all data points taken from the 1000-1500 ms post-stimulus window of the 900 or 1500 ms duration surrogate trials (the window was selected to allow a sensitive threshold which is not biased upward by the high power values of the onset response). Note that for the 900 ms trials the window falls in a presumably ‘off period’ whereas for the 1500 ms trials it represents an ‘on period’, so that the median value represents a reasonable cutoff between “active” and “baseline” power values. As in the duration-tracking analysis, trials which were too short to include a 1800 ms post-stimulus onset to onset window were excluded. The number of above-threshold data points was counted for each trial within a window of 600-1800 ms post-stimulus (minimizing the influence of the onset responses). The use of a count metric was selected from among several possible approaches to quantifying duration-dependence due to its relative insensitivity to amplitude differences, emphasizing instead differences in duration. To rule out the possibility that duration-dependent *offset* responses drive the correlation (e.g., longer offset response following longer stimulus duration), the number of above-threshold points in the 600 ms post-offset window for each trial was measured as well, and if it was positively correlated with stimulus duration its explained variance was regressed out prior to the main correlation analysis. Statistical significance of the correlations was derived parametrically from the correlation analysis, and corrected for multiple comparisons using FDR. Duration-dependence was assessed across both face and object stimulus trials for non-category-selective electrodes, and for the preferred category for category-selective electrodes.

### Comparison of Responses Across Single and Dual-Tasks

As a sensitive test for differences in the duration of the responses between the two tasks, we looked at 1500 ms stimulus duration trials and compared the number of above-threshold time points in each trial between the two tasks. The threshold for each electrode was set as the median of all data points taken from the 1000-1500 ms post-stimulus window of the 900 (where this window is post-offset) or 1500 ms duration trials (where the same window is pre-offset). As in the duration-dependence analysis the window was selected to allow a sensitive threshold which is not biased upward by the high power values of the onset response. The total number of above-threshold time points was counted for each trial and compared between tasks using a two-sample t-test. Results were corrected for multiple comparisons using FDR.

### Saccade Detection

Saccadic spike potentials detected from two temporal-pole electrodes were used to indicate the timing of eye movements in a single subject where spike potentials could be detected from electrodes just behind the eyes. Detection of the spike potentials was performed by high-pass filtering the signal from one of the two electrodes at 30 Hz, convolving it with a “canonical” saccadic spike model, and marking supra-threshold time points as saccadic spikes (Keren et al. 2010). To create the model, three healthy scalp EEG control subjects (female, age 26-31) participated in a scalp-EEG version of the same experiment with simultaneous eye-tracking (Eyelink 1000/2K, SR Research, Canada; binocular tracking at 500 Hz). Saccade rates were approximately 1.5/sec as predicted by the literature (Engbert 2006). Our spike potential model (Fig. 7B) was generated by high-pass filtering the radial electro-ocular channel of each EEG subject at 30Hz, and averaging it across saccade events and across subjects. After the convolution of the intracranial signal with the model, a 3 std. deviation threshold was set and any above-threshold activity was interpreted as a saccadic spike (Keren et al. 2010).

### Simulation of Saccade-Driven Sustained Activity

In addition to the analysis based on the saccadic spike potential, we used a simulation to investigate the hypothesis that sustained activity in the early visual cortex could be driven by successive transient visual-fixation-locked activity (that is, fixed-duration neural responses to the retinal update following a saccade, while the image is presented). We simulated a “best-case” scenario for such a hypothesis, consisting of a fixed-duration high-SNR response contingent on a saccade occurring while the stimulus is presented. We used saccade and microsaccade detection data from three healthy subjects (female, age 26-31) who participated in a scalp-EEG version of the same experiment with simultaneous eye-tracking. A high-SNR saccade-driven activity time course was simulated for each of the three subjects by convolving each occurrence of a saccade coinciding with stimulus presentation with a fixed-duration square response window. To avoid making any assumptions about the duration of the saccade-locked response, it was varied in separate simulations between 50 and 1000 ms in 50 ms steps. The first 500 ms of each trial were set as active by default to simulate a fixed stimulus-onset response. For each subject and simulated saccade-locked response duration, the overall simulated response durations were estimated across trials in the same way as in the duration-tracking analysis of the empirical data, and the mean absolute estimation error was recorded.

### Connectivity by Phase Coherence

In addition to local activity indexed by high-frequency broadband power, sustained perception could also be supported by duration-dependent long-range synchronization between cortical sites. We attempted to explore this hypothesis by looking at pairs of early visual and high-order cortical sites and test for stimulus-driven inter-electrode phase-locking, which is also sustained for the duration of the stimulus. We limited our analysis to three subjects who had both duration-tracking EVC electrodes and category-discriminating IT electrodes, resulting in a total of 10 EVC electrodes, 15 IT electrodes, and 52 same-brain electrode pairs. The small number of relevant trials made a full statistical analysis of duration-dependent phase-locking unfeasible, and thus we opted for a purely exploratory approach, simply testing for sustained phase coherence across trials for face stimuli with a 1500 ms duration, using the Inter-Trial Coherence (ITC) method (Makeig et al. 2004). The phase signal for each electrode was derived by using the Hilbert transform for the high-frequency band (30-200Hz). Statistical significance was determined with a 10,000-iteration randomization test whereby trials from one electrode were randomized. For all electrode pairs, the number of significant time points were approximately 5% for a p<0.05 threshold, indicating no ITC effect. Likewise, repeating the same analysis for all narrow frequency bands between 1 and 200 HZ (1 Hz steps) using wavelet analysis revealed no discernible effect apart from an initial low-frequency synchronization attributable to the evoked potential.

### Cross-Frequency Coupling

We also tested whether duration-dependent activity is manifested as stimulus-evoked cross-frequency coupling (CFC) within individual sites. We hypothesized that such coupling would occur between the phase of the dominant low-frequency oscillation and the amplitude of the broadband high frequency signal. As a first step, the dominant low-frequency peak at each visually-responsive electrode was determined by averaging the amplitude spectrum of all 1000 ms pre-stimulus segments to derive the smoothed baseline spectrum, fitting and subtracting the 1/f trend, and subsequently finding the maximal peak between 1 and 30 Hz. Phase amplitude coupling between the phase of the slow frequency and the amplitude of the high-frequency broadband (>30 Hz) signal was then computed across trials, to generate a time course of coupling magnitude as a function of time (Penny et al. 2008). This was done separately for 900 and 1500 ms duration trials. Since the effect of interest was not mere CFC but rather duration-dependent coupling, we compared the average coupling magnitude within the 1000-1500 ms window, between 900 ms and 1500 ms duration trials; if coupling is sustained for the duration of the stimulus, its mean magnitude in this window should be larger for the 1500 ms duration trials than for the 900 ones. Statistical significance was determined using a 10,000-iteration randomization test whereby for each iteration, the stimulus duration labels were shuffled and the resulting mean magnitude differences were used to generate a null-hypothesis distribution. Finally, the False Detection Rate method (Benjamini & Yosef 1995) was used to correct for multiple testing across electrodes. The analysis revealed no significant duration-dependent CFC.

## Author Contributions

Conceptualization, L.Y.D. and E.M.G.; Methodology, E.M.G. and L.Y.D.; Software, E.M.G. and T.G.; Formal Analysis, E.M.G.; Resources, R.T.K.; Writing – Original Draft, E.M.G., Writing – Review and Editing, L.Y.D., R.T.K. and T.G.; Visualization, E.M.G.; Supervision, L.Y.D. and R.T.K.; Funding Acquisition, L.Y.D. and R.T.K.

## Acknowledgments

The authors acknowledge Shlomit Yuval-Greenberg for preparing a prototype of this experiment and the stimuli, Josef Parvizi for facilitating ECOG data collection from his patients and for very helpful comments on the manuscript, and Rachel A. Kuperman, Kurtis I. Auguste and Edward F. Chang for making the collection of ECOG data from their patients possible. This work was supported by grant 2013070 from the US-Israel Binational Science Foundation (LYD and RTK), NINDS R37NS21135 (RTK), and the Nielsen Corporation (RTK).

## Supplemental Figures

**Figure S1, supplemental to Figure 3:**
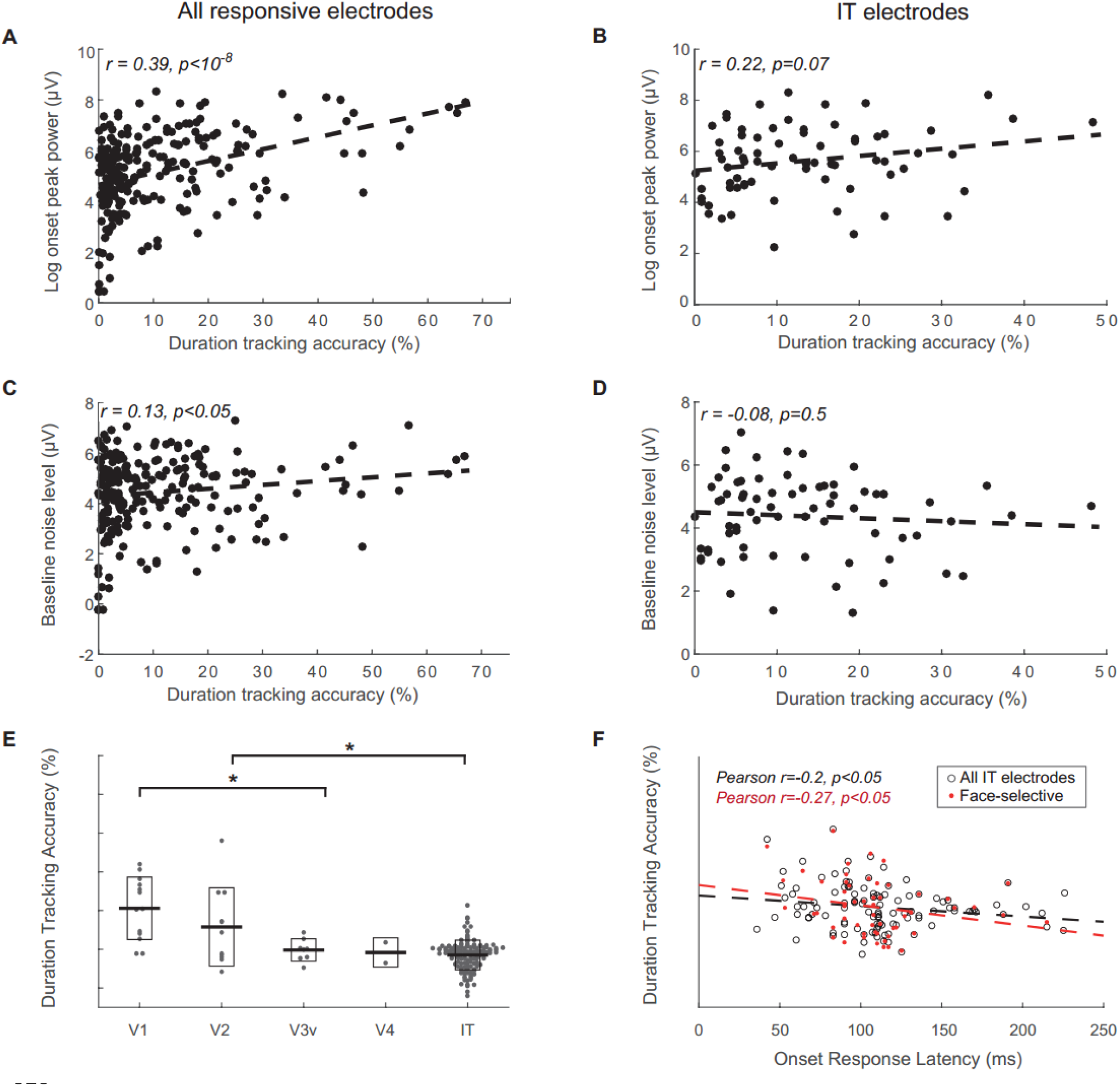
Controlling for baseline noise level and onset peak power. (**A,B**) the signal-to-noise ratio (SNR) of each electrodes’ HFB onset response, measured as the log of the ratio between the peak power of the 0-300 ms epoch in the average response for the preferred stimulus category (or for all categories for non-selective electrodes) and the standard deviation of the baseline, is correlated with duration-tracking accuracy across all right-hemisphere visually-responsive electrodes and across IT electrodes. (**C,D**) the baseline noise level for each electrode, measured as the log of the standard deviation of the baseline HFB power, is also mildly correlated with accuracy. Despite their correlation with duration-tracking accuracy, onset SNR and baseline noise level factors do not explain the negative correlation between duration-tracking accuracy and hierarchical position along the ventral stream. This was verified by regressing out the variance in the accuracy variable explained by these two factors using multiple regression, and analyzing the residual variance. As in the original analysis, accuracy was higher in EVC areas than in IT (t(166)=7.2, p<10^−5^), and in V1/V2 than V3v/V4 (t(32)=2.9, p<0.01; panel (E), compare to figure 3C), and within inferior temporal electrodes duration accuracy inversely correlated with response latency (panel F, compare to figure 3D). Correlating accuracy with position along the posterior-anterior axis produced comparable results (r=-0.31, p<0.01 for all IT electrodes, r=-0.36, p<0.01 for face-selective electrodes).

**Figure S2, supplemental to Figure 3:**
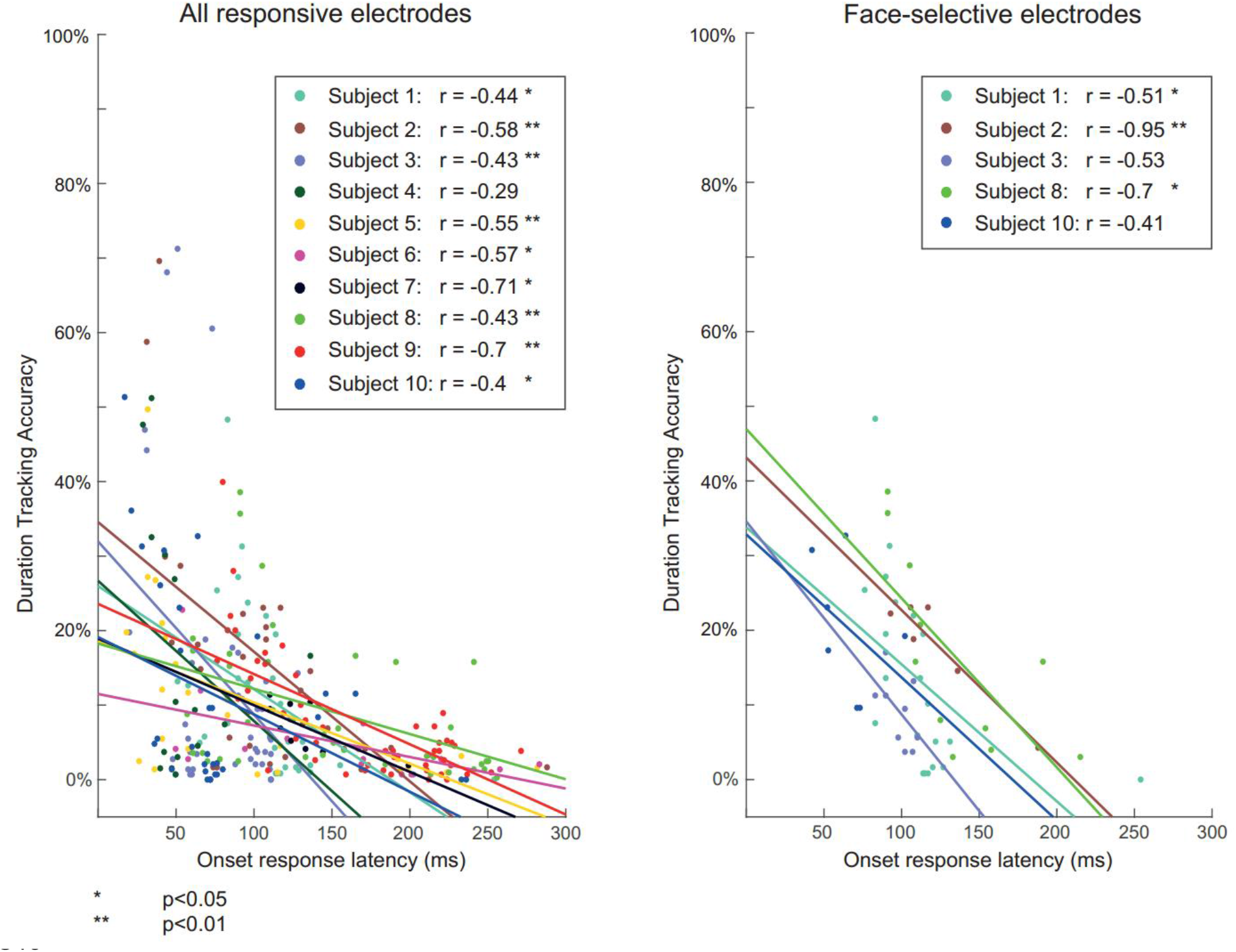
Single subject correlations. To verify that the negative gradient of duration-tracking accuracy is not driven by differences between subjects (i.e. to verify that tracking accuracy and position along the ventral stream are correlated within subjects, rather than that subjects with more posterior coverage have generally higher duration-tracking), we fitted a mixed-effect linear regression model to the single-trial duration-tracking data. The binomial result of each trial in each electrode (accurate/inaccurate response duration) was used as the dependent variable, and either the electrodes’ onset response latency, or its posterior-anterior position was used as an independent variable, with random intercepts and slopes for each subject and random intercepts for individual trials. The continuous measures of latency and anatomical position were used as the independent variables for the analysis of electrodes across all visual areas rather than region labels, because there were not sufficient data for the model to converge with an ordinal independent variable. The results indicated a significant fixed effect for both latency and position for all responsive electrodes, all right-hemisphere IT electrodes, all right-hemisphere face-selective electrodes, and all right-hemisphere object selective electrodes. Left panel: data points for all responsive electrodes divided by subject, and subject-specific regression lines. Right panel: same for right-hemisphere IT face-selective electrodes. Mixed-effects analysis was performed using R (www.r-project.org) with the lme4 software package.

**Figure S3, supplemental to Figure 3:**
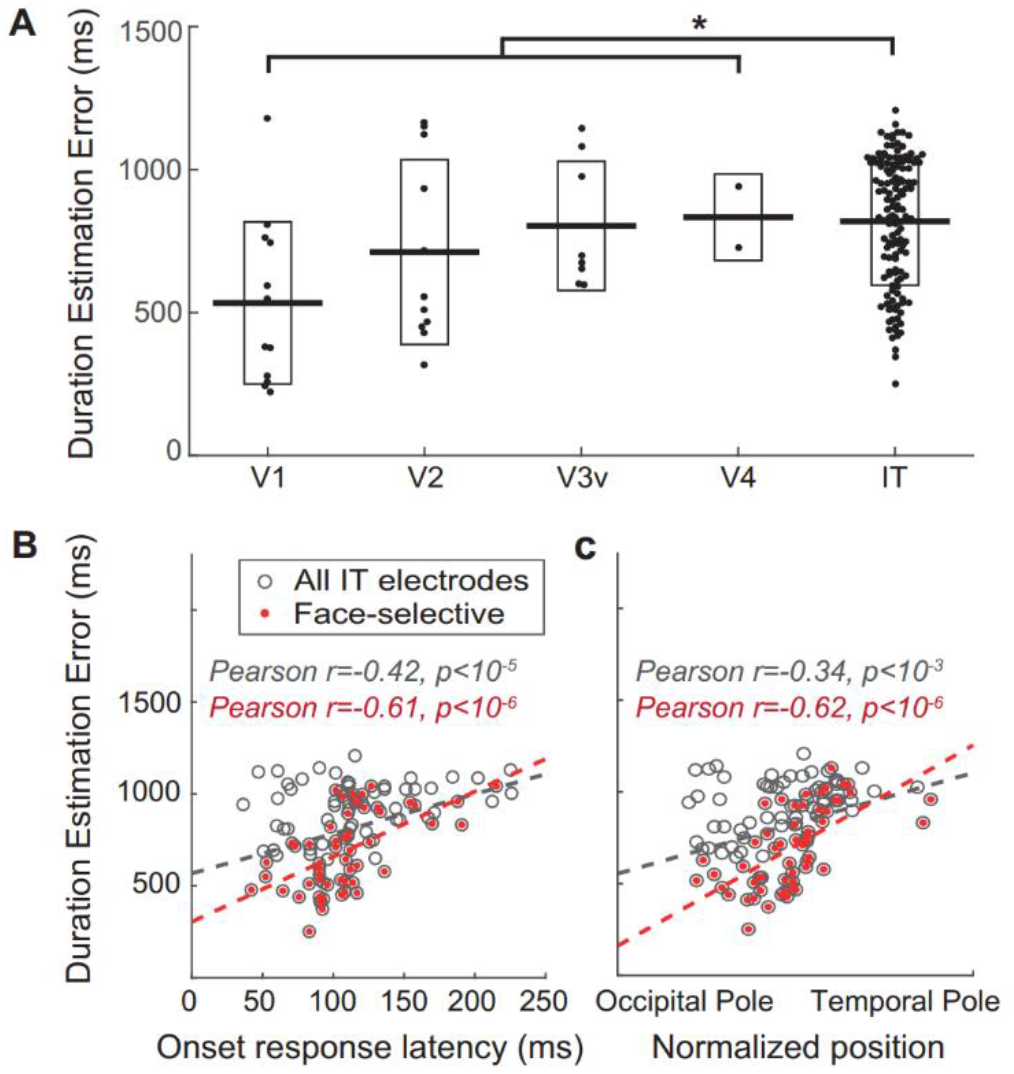
Duration tracking estimation error. To verify that the effects reported for duration-tracking accuracy were not dependent on the particular accuracy metric used, the same results reported in Fig. 3 were replicated using the mean duration estimation error as the dependent variable. (**A**) Relation between duration estimation error and hierarchical position along the ventral stream, based on a probabilistic atlas (EVC areas) and visual inspection (IT). Boxes correspond to standard deviation. Error in EVC areas is lower than in IT (t(166)=-3.2, p<0.01), and in V1/V2 compared to V3v/V4 (t(32)=-1.82, p<0.05). (**B**) duration-tracking within IT as a function of onset response latency as a proxy for hierarchical position along the ventral stream. (**C**) Same as (B), with hierarchical position measured as the electrode’s coordinate along the occipital-temporal axis.

**Figure S4, supplemental to Figure 3:**
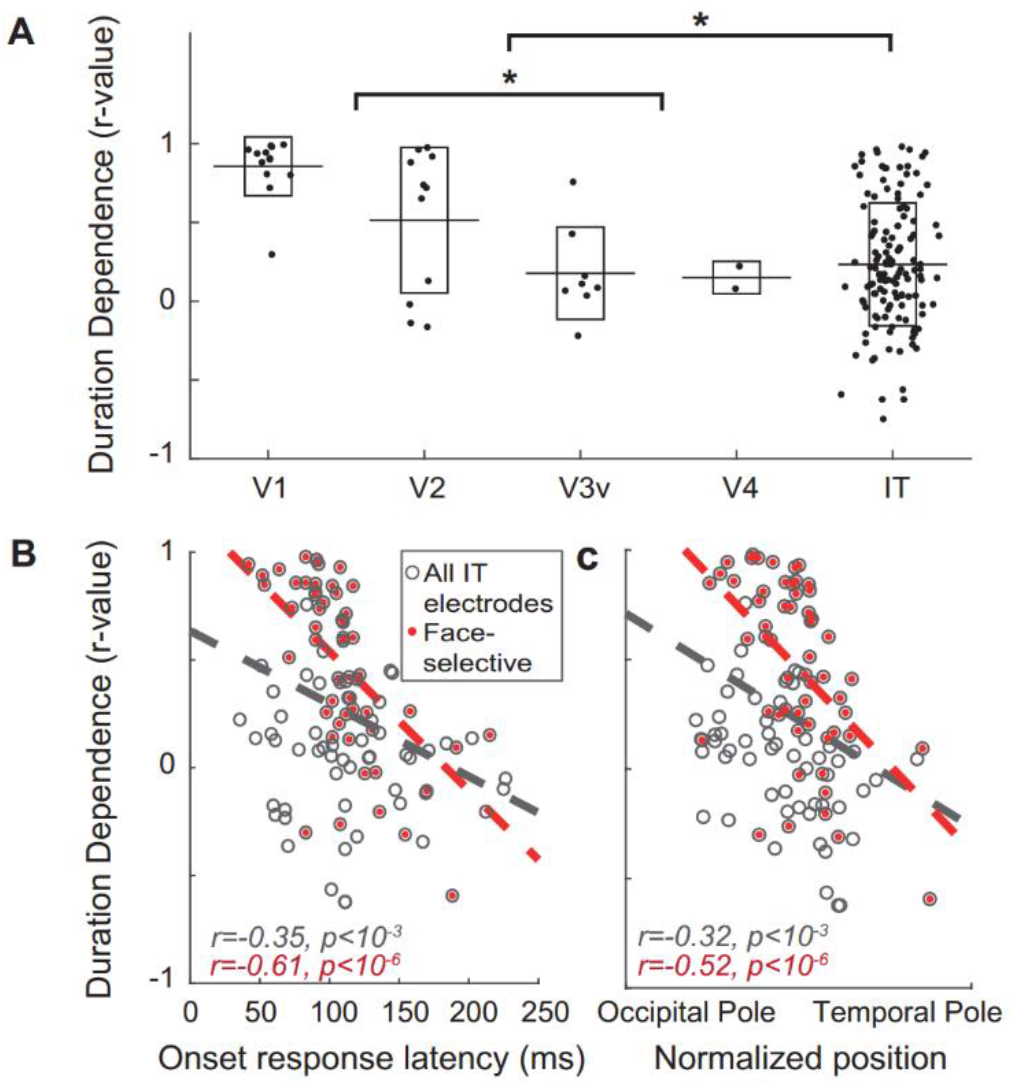
Decreasing duration dependence along the ventral stream. (**A**) The duration-dependence index (i.e., the correlation across trials between number of post-stimulus above-threshold time points and stimulus duration) as a function of region, plotted for all visually-responsive electrodes. Boxes indicate standard deviation. Duration dependence is higher in EVC areas than in IT (t(166)=4.08, p<10^−3^) and in V1/V2 than in V3v/V4 (t(32)=4.03, p<10^−3^). (**B**) Duration dependence within IT as a function of onset response latency. (**C**) Duration dependence within IT as a function of anatomical position along the posterior-anterior axis.

**Figure S5, supplemental to Figure 7:**
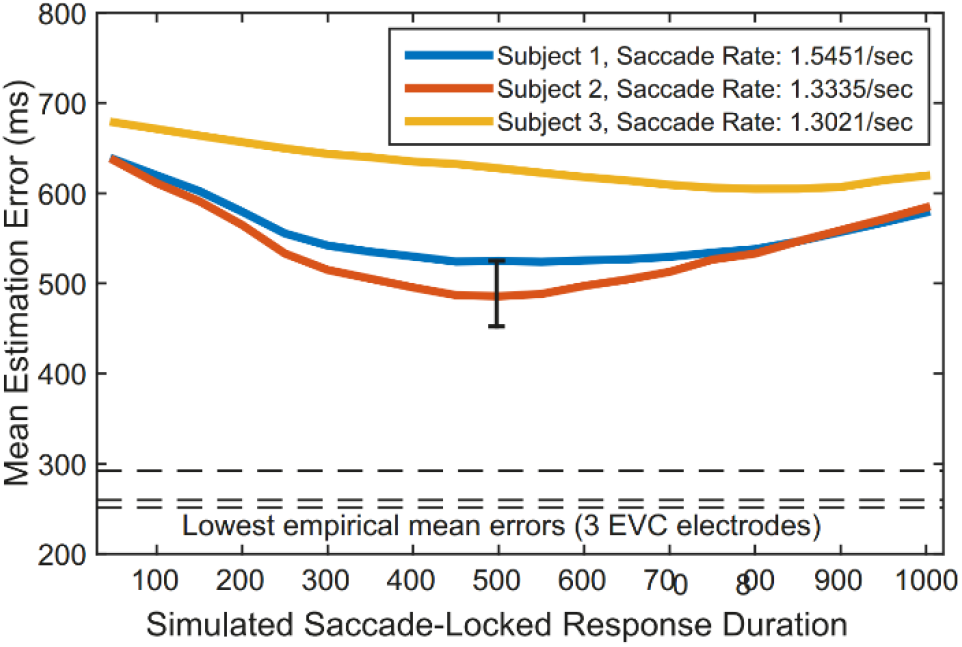
Simulations of duration dependent sustained activity driven by neural activity related to retinal shift following saccades. Mean error for stimulus duration estimation from simulated HFB power constructed based on individual saccades, as function of the simulated duration of the saccade evoked activity. Each line represents simulation based on saccades from one of three healthy subjects (see also figure 7). The lowest mean error achieved in the simulation was 485 ms, which is larger than that of 15 of the most strongly duration-dependent intracranial electrodes in the early visual cortex. The black error bar indicates 95% confidence interval across trials for the best-performing (lowest error) simulation, the three lowest errors from intracranial electrodes are shown as horizontal broken lines). We conclude that it is unlikely that the robust duration-tracking responses in early visual cortex were the result of summed transient responses driven by eye movements, regardless of the duration of these saccade-related bursts.

**Figure S6, supplemental to Figure 3:**
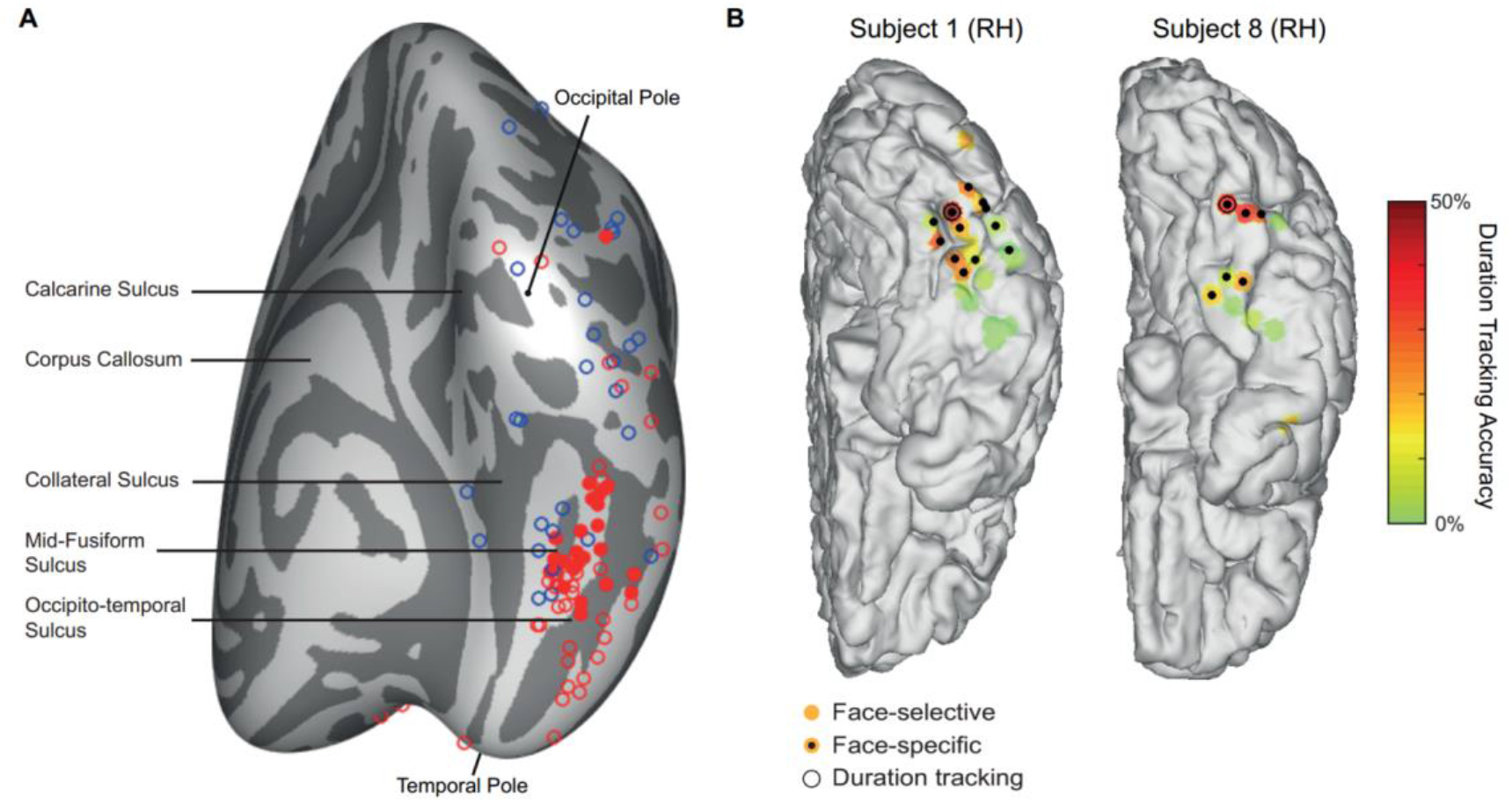
Category specificity. (**A**) Category-specificity to faces, objects and animals was assessed by a conjunction of statistically significant selectivity of an electrode to one category relative to each of the two other categories (e.g. face-specificity entails {face>object & face>animal}). Empty red and blue circles correspond to face-selective and object-selective electrodes, respectively. Filled red circles correspond to face-specific electrodes. There were no object-specific or animal-specific electrodes. The electrodes are shown on template brain (FreeSurfer average brain), from a medial-posterior-inferior aspect. Dark regions indicate sulci. (**B**) Face-selective vs. face-specific electrodes in two individual brains from the subjects shown in Fig. 5 who had face-selective category-tracking electrodes. Colored patches indicate face selective electrodes; a black dot indicates that the response is also face-specific. The black circles indicate significant duration-tracking. The distinction between strongly duration-tracking posterior sites and weakly duration-tracking anterior sites can be seen for this subset of face-specific electrodes.

## Footnotes

* The fact that category selectivity seems to emerge rather than decrease in downstream sites does not contradict this principle, as category information is fully present in the retina, and therefore it is not gained, but rather made explicit by the integration of low-level visual features as well as stored templates into representations which are increasingly divergent (e.g. faces vs. non-faces) and invariant to low-level detail (e.g. to stimulus contrast, Avidan et al. 2002).

